# Cost-Effective Purification of Endotoxin-Free LIF, IL-2, and IL-33

**DOI:** 10.64898/2026.05.29.728897

**Authors:** Jae-Yoon Kim, Jessica Xin, I-Chih Kuo, Si Xuan Chen, Ahmed Kabil, Kai-Wei Chang, Nika Shakiba, Kelly M. McNagny, Yanpu He

## Abstract

We describe a cost-effective *Escherichia coli* (*E. coli*)-based platform for producing endotoxin-free cytokines. As a proof of concept, we applied this protocol to purify four representative human and mouse cytokines – mouse leukemia inhibitory factor (mLIF), human interleukin-2 (hIL-2), and human and mouse interleukin-33 (hIL-33 and mIL-33) – and demonstrated their bioactivity to be equivalent to commercial counterparts for direct use in stem cell culture and immune cell activation, both *in vitro* and *in vivo*. Reagent costs for producing these proteins are approximately 5%-10% of commercial list prices. This platform is readily adaptable to other costly cytokines and growth factors, providing a scalable and affordable approach to accelerate research in cell biology, tissue engineering, and biomanufacturing.

## Before you begin

Cytokines are indispensable reagents in cell biology research, where they are routinely used to modulate cell function, direct proliferation and differentiation^1^. Despite their widespread use, access to high-quality recombinant cytokines remains a major practical bottleneck. Many cytokines are costly to purchase (approximately $1,000-$8,000 CAD per mg from manufacturers like *STEMCELL Technologies* and *Thermo Fisher Scientific*) and in-house produced recombinant proteins may contain endotoxin or impurities that confound sensitive *in vitro* and *in vivo* studies^2^. These limitations restrict experimental scales and reproducibility.

Here, we report the development of a simple and robust *Escherichia coli* (*E. coli*)-based platform for producing clean, biologically active cytokines at substantially reduced cost. To evaluate the generalizability of this approach, we applied the platform to four representative cytokines spanning both stem cell and immunology applications: mouse leukemia inhibitory factor (mLIF), human interleukin-2 (hIL-2), and human and mouse interleukin-33 (hIL-33 and mIL-33). These cytokines vary in size, structure, species, and biological function, providing a stringent test of the platform’s versatility.

## Innovation Statement

Our platform integrates three key features into a single workflow. First, we employ ClearColi™ *E. coli*, an engineered endotoxin-free strain, to eliminate lipopolysaccharide contamination at the source^3^. Proteins purified from ClearColi™ have been reported to exhibit approximately 3.6 × 10^2^ endotoxin units (EU)/mL, approximately 97% reduction in limulus amebocyte lysate (LAL) activity compared to proteins purified from BL21 strains^4^. Second, to enable efficient expression of cytokines containing rare mammalian codons, we supplement ClearColi™ with rare codon tRNAs via the pRARE plasmid, a strategy adapted from the Rosetta expression system^5^. Third, we implement a cost-efficient expression tag removal approach in which maltose-binding protein (MBP) tags^6^ are cleaved directly in crude bacterial lysates using tobacco etch virus (TEV) and human rhinovirus 3C (HRV3C) proteases.Combining crude bacterial lysates enables single-step MBP tag cleavage, avoiding the multiple immobilized metal affinity purification and protease processing steps required in traditional workflows. This is completed prior to immobilized metal ion affinity pull-down, thereby reducing processing time and avoiding yield loss. While this workflow was optimized for MBP-His-tagged cytokines, alternative constructs or solubility tags may require further optimization of protease ratios and downstream purification conditions.

Using this strategy, all four cytokines were purified to high purity with minimal endotoxin contamination, retained biological activity comparable to commercially available reagents, and could be produced at a cost of 5%-10% of commercial list prices of these proteins (see Table 1 for detailed cost breakdown).

**Table 1.**
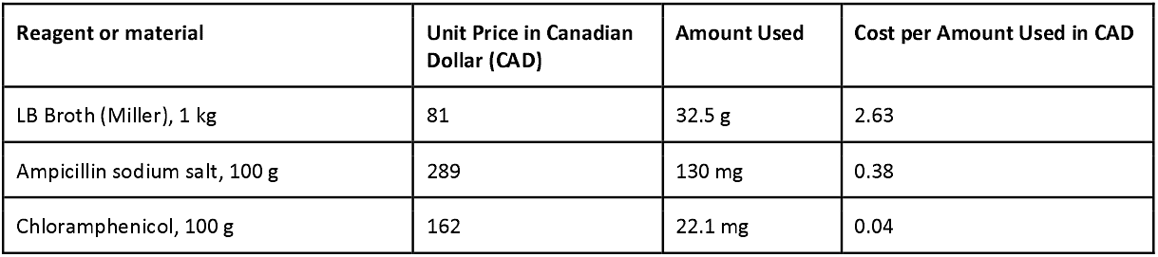
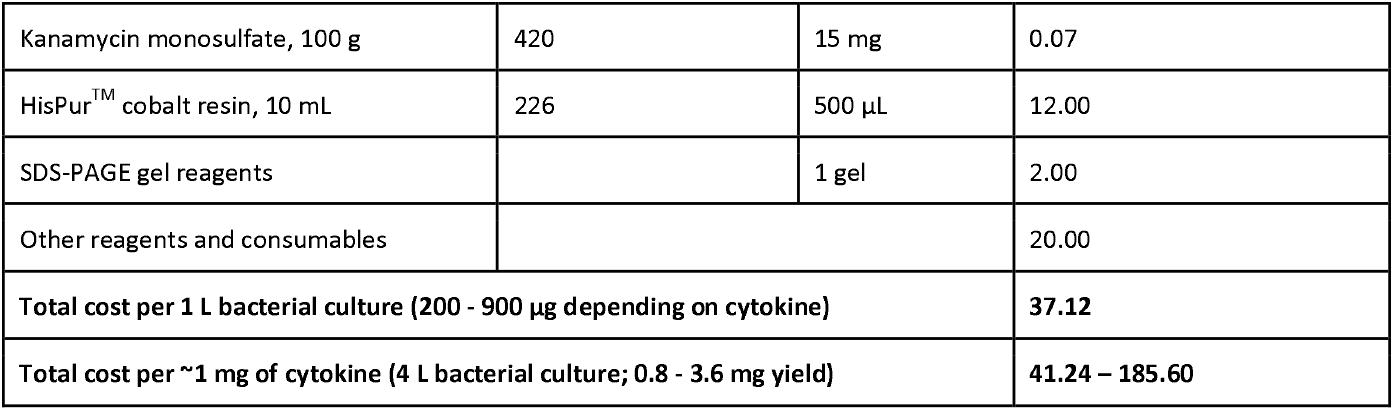
Estimated reagent and consumable costs for cytokine production.

| Reagent or material | Unit Price in Canadian Dollar (CAD) | Amount Used | Cost per Amount Used in CAD |
| --- | --- | --- | --- |
| LB Broth (Miller), 1 kg | 81 | 32.5 g | 2.63 |
| Ampicillin sodium salt, 100 g | 289 | 130 mg | 0.38 |
| Chloramphenicol, 100 g | 162 | 22.1 mg | 0.04 |
| Kanamycin monosulfate, 100 g | 420 | 15 mg | 0.07 |
| HisPur™ cobalt resin, 10 mL | 226 | 500 µL | 12.00 |
| SDS-PAGE gel reagents |  | 1 gel | 2.00 |
| Other reagents and consumables |  |  | 20.00 |
| Total cost per 1 L bacterial culture (200 - 900 µg depending on cytokine) |  |  | 37.12 |
| Total cost per ~1 mg of cytokine (4 L bacterial culture; 0.8 - 3.6 mg yield) |  |  | 41.24 – 185.60 |

**Note:** Costs are estimated based on single-use reagent consumption per 1 L bacterial culture and may vary depending on supplier and scale.

### Institutional permissions

All mouse experiments were performed in accordance with protocols approved by the University of British Columbia Animal Care Committee and followed guidelines established by the Canadian Council on Animal Care. The *in vivo* mouse IL-33 challenge was approved by the University of British Columbia’s Biosafety Committee through protocol A24-0223: Microbiota and dietary metabolites in atopy and inflammation.

Human peripheral blood samples were collected under protocols approved by the University of British Columbia Research Ethics Board. Informed consent was obtained from all donors prior to sample collection.

The use of human peripheral blood mononuclear cells (PBMCs) was approved by the University of British Columbia’s Biosafety Committee through ethics protocol H23-03468: Characterizing Immune Cell Subsets in ILD.

### Preparing LB broth (Miller)

**Timing: 90 minutes**

1. Completely dissolve 25 g of LB broth (Miller) per 1 L of Milli-Q water.
2. Autoclave the solution at 121°C for 20 min and allow it to cool to room temperature.
3. Add appropriate antibiotics as required (working concentrations: 100 mg/L ampicillin, 17 mg/L chloramphenicol, and/or 50 mg/L kanamycin monosulfate).
4. Store the solution at 4°C for long-term use.

### Preparing LB broth (Miller) agar plates

**Timing: 90 minutes**

1. Completely dissolve 25 g of LB broth (Miller) and 12 g of agar per 1 L of Milli-Q water.
2. Autoclave the solution at 121°C for 20 min and allow it to cool to approximately 50°C.
3. Add appropriate antibiotics as required (final concentrations: 100 mg/L ampicillin, 17 mg/L chloramphenicol, and/or 50 mg/L kanamycin monosulfate).
4. Gently mix and pour approximately 20 mL per plate (approximately 400 mL total into 20 Petri dishes).
5. Leave plates at room temperature overnight to solidify and dry.
6. Store plates at 4°C for long-term use.

### Preparing chemically competent pRARE-containing ClearColi™

**Timing: 2 days**

1. Extract the rare codon tRNA plasmid (pRARE) from Rosetta DE3 competent cells using the Monarch® Spin Plasmid Miniprep Kit according to the <u>manufacturer’s instructions</u>. **Note:** While Monarch® Spin Plasmid Miniprep Kit was used for plasmid extraction in this protocol, other available plasmid extraction kits are suitable for use.
2. Transform pRARE into ClearColi™ BL21(DE3) electrocompetent cells via electroporation using the Biorad Gene Pulser Xcell Microbial System.
  a. Mix 45 μL of ClearColi™ BL21(DE3) electrocompetent cells with 5 μL of pRARE plasmid in a 0.1 cm gap electroporation cuvette.
  b. Electroporate using the following settings: 180 V, 25 μF, and 200 Ω. **Note:** The electroporation time constant should be approximately 5 ms.
3. Plate 50 μL of transformed cells onto a chloramphenicol agar plate and incubate overnight at 37°C. **Note:** According to the manufacturer, the transformation efficiency of ClearColi™ BL21(DE3) electrocompetent cells is 1 x 10^9^ colony forming units (cfu)/µg of plasmid^7^, which was consistent with our observation.
4. Pick a single colony and inoculate into 5 mL LB broth (Miller) supplemented with 17 μg/mL chloramphenicol. Incubate overnight at 37°C with shaking at 220 rpm.
5. Dilute the overnight culture into 40 mL LB broth (Miller) containing 17 μg/mL chloramphenicol and grow at 37°C with shaking (220 rpm) for 4 - 6 hr, until OD_600_ reaches 0.5–0.6.
6. Place the culture on ice for 15 min. **CRITICAL:** Keep cells on ice at all times after reaching mid-log phase to maintain competency.
7. Centrifuge at 4,000 x g for 5 min.
  a. Discard the supernatant.
  b. Resuspend the pellet in 10 mL ice-cold Inoue buffer.
  c. Repeat the wash step a total of four times.
  d. On the final resuspension, resuspend in 800 μL Inoue buffer and add 60 μL DMSO.
8. Aliquot 100 μL per tube, flash freeze in liquid nitrogen, and store at −80°C.

**Note:** Competent cells should be stored at −80°C and can be used long term. The transformation efficiency will decrease after every freeze-thaw cycle.

### Transforming pRARE-containing ClearColi™ with cytokine or protease expression plasmid

**Timing: 2 days**

1. Transform the cytokine or protease plasmid into chemically competent pRARE-containing ClearColi™ via heat shock.
  a. Mix 45 μL of chemically competent pRARE-containing ClearColi ™ with 5 μL of cytokine or protease plasmid.
  b. Incubate the mixture on ice for 30 min.
  c. Perform heat shock at 42°C for 30 seconds in a water bath.
  d. Incubate the mixture on ice for 5 min.
  e. Add 100 μL of SOC media to the mixture and incubate at 37°C for 1 hr.
2. Plate 50 μL of transformed cells onto a chloramphenicol and ampicillin agar plate and incubate overnight at 37°C. **Note:** The transformation efficiency for competent cells prepared using the Inoue method is typically between 1 × 10^8^ to 3 × 10^8^ colony forming units (cfu)/µg of plasmid ^8^, which was consistent with our observations.
3. Pick a single colony and inoculate into 5 mL LB broth (Miller) supplemented with 17 μg/mL chloramphenicol and 100 μg/mL ampicillin. Incubate overnight at 37°C with shaking at 220 rpm.
4. Prepare a bacterial glycerol stock for long-term use.
  a. Mix 100 μL of 50% glycerol in water with 400 μL of the overnight bacterial culture, and store in −80°C freezer.

### Preparing a 15% SDS-PAGE gel

**Timing: 90 minutes**

1. Prepare a 15% sodium dodecyl sulfate (SDS)-polyacrylamide gel using the Bio-Rad Mini-PROTEAN hand-cast system. After assembling the glass plates, perform a leak test with water to ensure it is well sealed.
  a. Resolving gel: Mix 2.5 mL of 30% acrylamide, 1.25 mL of 1.5 M Tris-HCl (pH 8.8), 50 μL of 10% (w/v) SDS in ddH_2_O, 1.13 mL ddH_2_O, 2.5 μL Tetramethylethylenediamine (TEMED), and 25 μL of 10% (w/v) ammonium persulfate (APS) in ddH_2_O. Add approximately 4.5 mL in between the glass plates.
  b. Overlay with 1 mL of 100% ethanol to flatten the resolving gel.
  c. Wait 45 minutes for the gel to be fully polymerized, decant ethanol after polymerization.
  d. Stacking gel: Mix 198 μL of 30% acrylamide, 378 μL of 0.5 M Tris-HCl (pH 6.8), 15 μL of 10% SDS, 900 μL ddH_2_O, 1.5 μL TEMED, and 7.5 μL 10% APS. Pour on top and insert comb. Wait 30 minutes for the gel to be fully polymerized.
2. Store gels at 4°C until use. **CRITICAL:** 10% APS needs to be prepared fresh weekly and stored at 4°C. Ensure 10% APS and TEMED are added last to initiate polymerization and work quickly after mixing.

**Key resources table**

| REAGENT or RESOURCE | SOURCE | CATALOG # |
| --- | --- | --- |
| **Bacterial Strains** |  |  |
| ClearColi<sup>TM</sup> BL21(DE3) electrocompetent cells | Biosearch Technologies | 60810 |
| BL21(DE3) competent *E. coli* | New England Biolabs | C2527I |
| NEB<sup>®</sup> Stable Competent *E. coli* (High Efficiency) | New England Biolabs | C3040I |
| Rosetta(DE3) Competent Cells | Novagen | 70954-3 |
| **Recombinant DNA** |  |  |
| pMAL-7xHis-TEV-hexa | Addgene | 222868 |
| pET-NT*-HRV3CP | Addgene | 162795 |
| IL-2 pMalLP | Addgene | 41808 |
| **Chemicals, reagents, and antibiotics** |  |  |
| LB broth (Miller) | BioShop | LBL407.1 |
| Agar | BioShop | AGR003.1 |
| Ampicillin sodium salt | BioShop | AMP201.100 |
| Chloramphenicol | BioShop | CLR201.100 |
| Kanamycin monosulfate | BioShop | KAN201.100 |
| Isopropyl-D-1-thiogalactopyranoside (IPTG) | BioShop | IPT002.50 |
| Imidazole | BioShop | IMD510.500 |
| Sodium phosphate dibasic, heptahydrate | BioShop | SPD579.5 |
| Sodium phosphate monobasic | BioShop | SPM306.5 |
| Sodium chloride | BioShop | SOD004.1 |
| Triton X-100 | Millipore Sigma | X100-100ML |
| Lysozyme | BioShop | LYS702 |
| Phenylmethanesulfonyl fluoride (PMSF) | BioShop | PMS444.100 |
| Dithiothreitol (DTT) | BioShop | DTT001.25 |
| HEPES | BioShop | HEP005.1 |
| β-mercaptoethanol | BioShop | MER002.500 |
| Sodium dodecyl sulfate (SDS) | BioShop | SDS003.500 |
| Tris base | BioShop | TRS001.1 |
| 10X Tris, Glycine, SDS | BioShop | TGS222.4 |
| Tris | BioShop | TRS003.1 |
| Glycerol | BioShop | GLY001.1 |
| Coomassie Brilliant Blue G-250 | BioShop | CBB555.100 |
| Methanol | BioShop | MET302.4 |
| Acetic acid | BioShop | ACE222.4 |
| PBS tablets | BioShop | PBS404.500 |
| Bovine serum albumin (BSA) | BioShop | ALB007.500 |
| Bromophenol Blue G-250 | BioShop | BRO777.50 |
| Dimethyl sulfoxide (DMSO) | BioShop | DMS555.500 |
| Manganese chloride, tetrahydrate | BioShop | MAN222.500 |
| Piperazine-N,N'-bis(2-ethanesulfonic acid) (PIPES) | BioShop | PIP777.500 |
| Potassium chloride | BioShop | POC999.1 |
| 30% Acrylamide/Bis-acrylamide | BioShop | ACR010.500 |
| Tetramethylethylenediamine (TEMED) | BioShop | TEM001.50 |
| Ammonium persulfate (APS) | BioShop | AMP001.100 |
| Iso-propyl alcohol (IPA) | BioShop | ISO920.4 |
| IRIS11 Prestained protein ladder | FroggaBio | PMI11-0500 |
| **Commercial assays, kits, and reagents** |  |  |
| HisPur™ cobalt resin | Thermo Fisher | 89964 |
| Pierce™ BCA Protein Assay Kit | Thermo Fisher | 23225 |
| Zeba™ Spin Desalting Columns | Thermo Fisher | A40002469 |
| Monarch® Spin Plasmid Miniprep Kit | New England Biolabs | T1110L |
| **Equipment** |  |  |
| Fast protein liquid chromatography (NGC Quest 10 Plus) | Bio-Rad | 7880003 |
| ENrich™ SEC70 10 x 300 Column | Bio-Rad | 7801070 |
| NGC Fraction Collector | Bio-Rad | 17002070 |
| Gene Pulser Xcell Microbial System | Bio-Rad | 1652662 |
| Gene Pulser/MicroPulser Electroporation Cuvettes, 0.1 cm gap | Bio-Rad | 1652083 |
| Avanti J-26s XP High-Speed Centrifuge | Beckman Coulter | B14535 |
| JLA-8.1000 rotor | Beckman Coulter | 363688 |
| Q125 Sonicator | QSONICA | Q125-110 |

**Note:** The key resources and materials listed above apply specifically to the protein purification workflow. Resources and materials required for bioactivity assays are provided in *Tables S2–S6*.

## Materials and equipment

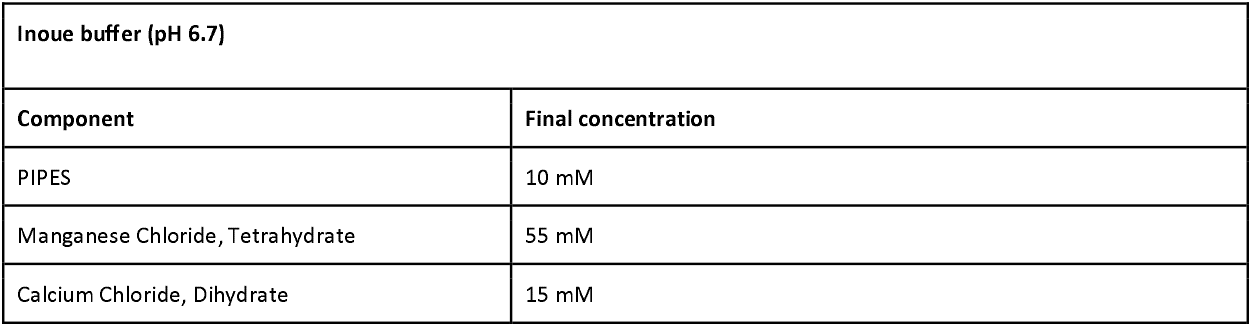
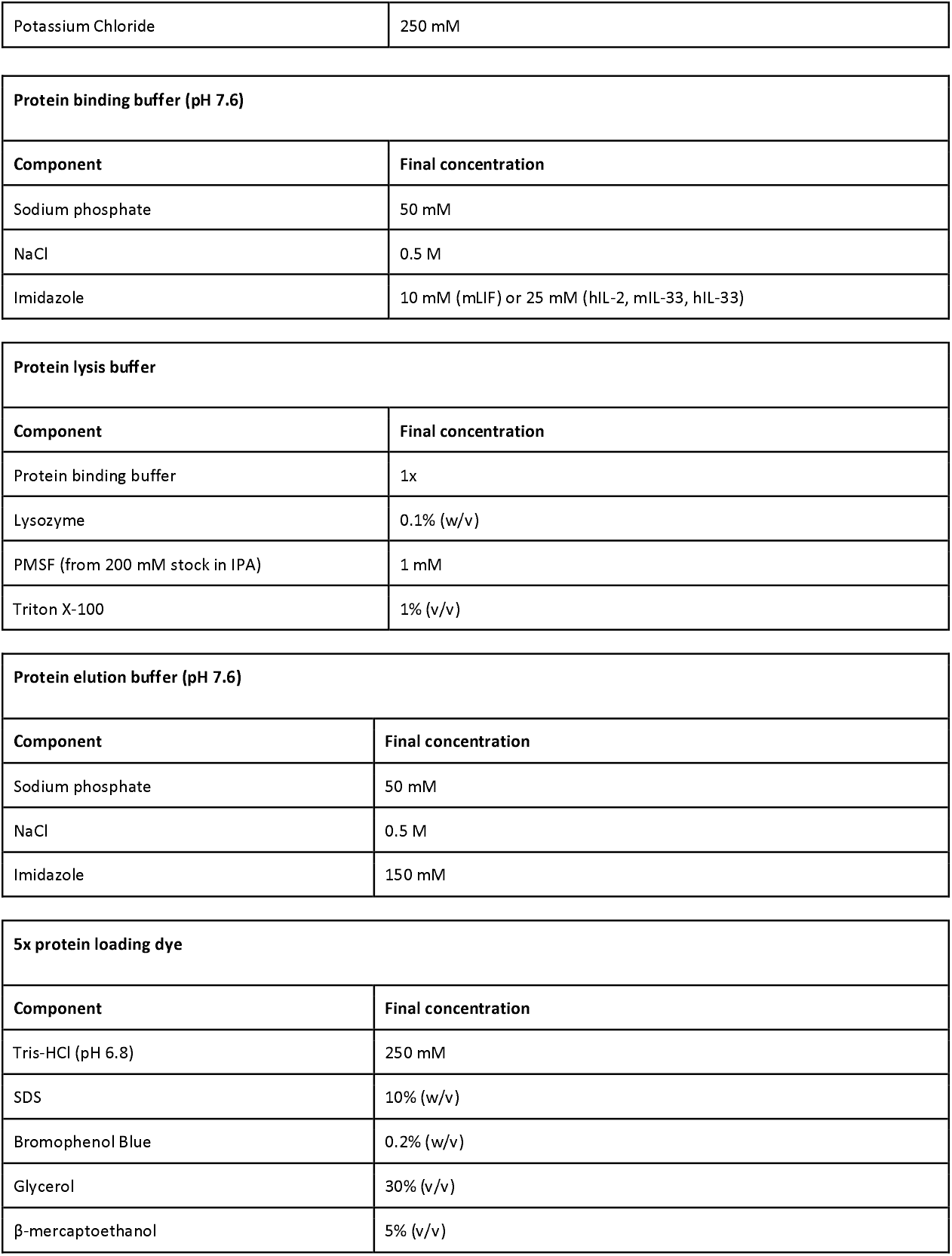
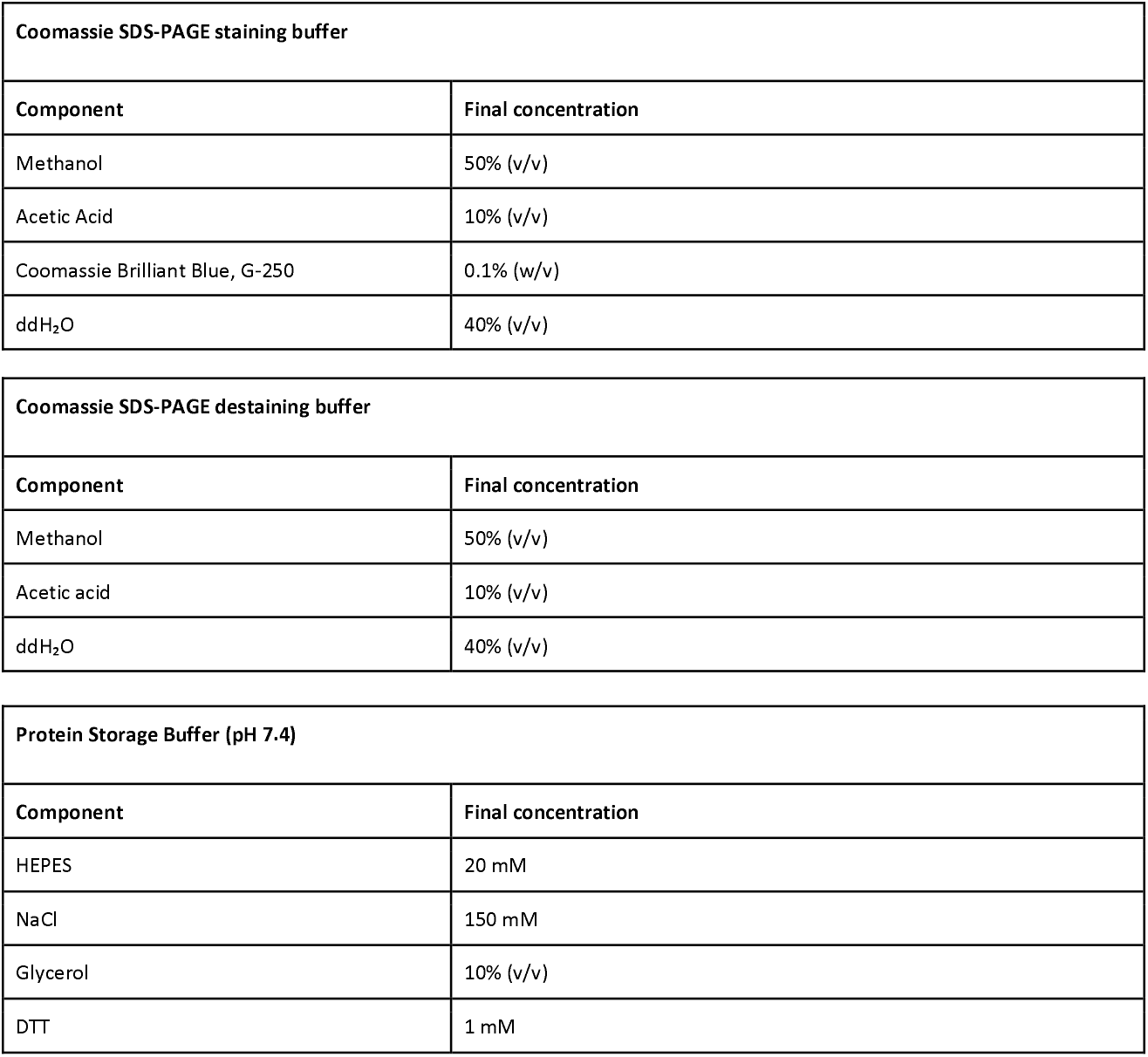

### Step-by-step method details

### Protein Purification

### Growing the Bacterial Culture

**Timing: 1 day (for step 1-2)**

This step cultures ClearColi™ cultures containing pRARE and the cytokine or protease plasmid of interest. Plasmid sequences can be found in **Table S1**.

1. Inoculate bacteria containing the cytokine plasmid from glycerol stock into 40 mL LB broth (Miller) supplemented with 17 μg/mL chloramphenicol and 100 μg/mL ampicillin. Incubate overnight at 37°C while shaking at 220 rpm. Troubleshooting <u>problem 1</u>.
2. Inoculate bacteria containing the protease plasmid from glycerol stock into 40 mL LB broth (Miller) supplemented with 17 μg/mL chloramphenicol and either 100 μg/mL ampicillin or 50 μg/mL kanamycin monosulfate. Incubate overnight at 37°C while shaking at 220 rpm. Troubleshooting <u>problem 1</u>.

**Note:** All cytokine expression vectors carry ampicillin resistance, and the pRARE plasmid carries chloramphenicol resistance. The TEV protease plasmid carries an ampicillin resistance gene, whereas the HRV 3C protease plasmid carries a kanamycin resistance gene.

### Inducing Protein Expression in *E. coli*

**Timing: 2 days (steps 3–7)**

This step induces recombinant protein expression using IPTG.

3. Dilute the 40 mL overnight culture containing the cytokine plasmid into 1 L LB broth (Miller) supplemented with 17 μg/mL chloramphenicol and 100 μg/mL ampicillin. Grow at 37°C while shaking at 180 rpm until OD_600_ reaches 0.4-0.7.
4. Dilute the 40 mL overnight culture containing the protease plasmid into 300 mL LB broth (Miller) supplemented with 17 μg/mL chloramphenicol and either 100 μg/mL ampicillin or 50 μg/mL kanamycin monosulfate. Grow under the same conditions until OD_600_ reaches 0.4–0.7.
5. Place cultures on ice for 15 min.
6. Induce protein expression by adding IPTG to a final concentration of 0.5 mM.
7. Incubate cultures at 16°C while shaking at 180 rpm for 48 h.

### Releasing and Purifying Cytokines from *E. coli*

**Timing: ~10 hours**

This step lyses bacterial cells and purifies the cytokine of interest as illustrated in *Figure 1. Note*: This protocol is optimized for 1 L bacterial culture. Volumes can be scaled linearly.

8. Centrifuge cultures at 4,000 x g for 20 min using an Avanti J26S XP centrifuge with JLA-8.1 rotor.
  a. Discard supernatant and resuspend pellet in 30 mL PBS.
  b. Transfer to a 50 mL conical tube.
9. Centrifuge at 4,000 x g for 15 min.
  a. Discard supernatant and resuspend pellet in 20 mL protein lysis buffer.
10. Incubate lysate at room temperature for 20 min while shaking at 120 rpm. **Note:** The lysate will appear viscous due to genomic DNA release.
11. Place lysate on ice-water and perform probe sonication via Q125 Sonicator using parameters listed in Table 2 to further lyse cells and shear DNA. **Note:** After sonication, the lysate should appear darker and less viscous.
12. Centrifuge lysate at 4°C and 4,000 x g for 10 min.
  a. Transfer supernatant to a clean tube.
  b. Discard pellet. **Note:** The pellet can be saved for troubleshooting purposes. Troubleshooting problem 3.
13. Combine TEV protease lysate with MBP-fused mLIF lysate at a 1:7 volume ratio, which we found to be the optimal ratio to minimize TEV carry over into the FPLC purification. Incubate overnight on ice while shaking at 120 rpm. (For hIL-2, mIL-33, and hIL-33, combine HRV 3C protease lysate with MBP-fused hIL-2, mIL-33, or hIL-33 lysate at a 1:15 volume ratio^9^. Incubate on ice for 1 hr while shaking at 120 rpm.) **Note:** For optimal protease activity, proteases and cytokines should be expressed and purified on the same day. Based on optimization experiments, HRV 3C protease exhibited higher cleavage efficiency than TEV protease, requiring 1 hr incubation for maximal cleavage, whereas TEV protease required overnight incubation.
14. Add 500 μL HisPur™ cobalt resin to the lysate and incubate on ice for 1 hr while shaking at 120 rpm.
15. Centrifuge at 4 x g for 2 min.
  a. Discard supernatant.
16. Wash resin with 1 mL protein binding buffer.
  a. Centrifuge at 4 x g for 2 min and discard supernatant, gently aspirate the supernatant to avoid removing cobalt resin.
  b. Repeat wash 3 times. **Note:** The imidazole concentration in the protein binding buffer is determined experimentally and will vary for different cytokines. The optimal imidazole concentration can be determined by testing multiple concentrations and observing which imidazole concentration will retain high protein yield with minimal impurities on SDS-PAGE gels. Refer to problem 2.
17. Elute cytokine by resuspending resin in 1.5 mL protein elution buffer.
  a. Centrifuge at 4 x g for 2 min.
18. Transfer 1.5 mL eluted cytokine to a clean tube on ice.
19. Filter eluted cytokine using a 1 mL syringe a sterile hydrophilic syringe filter with 0.2 μm porosity.
20. Perform size exclusion chromatography using the Bio-Rad NGC Quest Plus system with ENrich SEC70 column and fraction collector using parameters listed in Table 3. Chromatography run plots for each cytokine is shown in **Figure S1**. **Note:** When purifying a protein for the first time with the size exclusion column, collect a broader range of fractions to avoid losing the target product into the waste stream.
21. Determine protein concentration using the Pierce™ BCA Protein Assay Kit according to the <u>manufacturer’s instructions</u>. **Note:** While Pierce™ BCA Protein Assay Kit was used to determine protein concentration in this protocol, other protein assays (ie. Bradford assay) are suitable for use.
22. Analyze purified cytokines by 15% SDS-PAGE as illustrated in **Figure 1**. Troubleshooting <u>problem 2</u> and <u>problem 3</u>.
  a. Load 20 μL of protein sample with SDS loading buffer and heated at at 95°C for 5 mins, into each lane of the SDS-PAGE gel.
  b. Run SDS-PAGE using the following settings: 180 V, 500 mA, and 1 hr.
  c. Stain the gel in Coomassie SDS-PAGE staining buffer for 15 min at room temperature with shaking at 110 rpm.
  d. Destain the gel in Coomassie SDS-PAGE destaining buffer for 1hr at room temperature with shaking at 110 rpm.
  e. Wash the gel in water at room temperature overnight with shaking at 110 rpm, the gel will then be ready for imaging.
23. Perform buffer exchange to protein storage buffer using Zeba™ Spin Desalting Columns according to the manufacturer’s instructions.
24. Aliquot 100 μL per tube, and store at −80°C. Troubleshooting problem 4. Note: Cytokines can be stored in PBS at −20°C for short-term use. For long-term storage, cytokines should be stored in protein storage buffer at −80°C.

**Figure 1.**
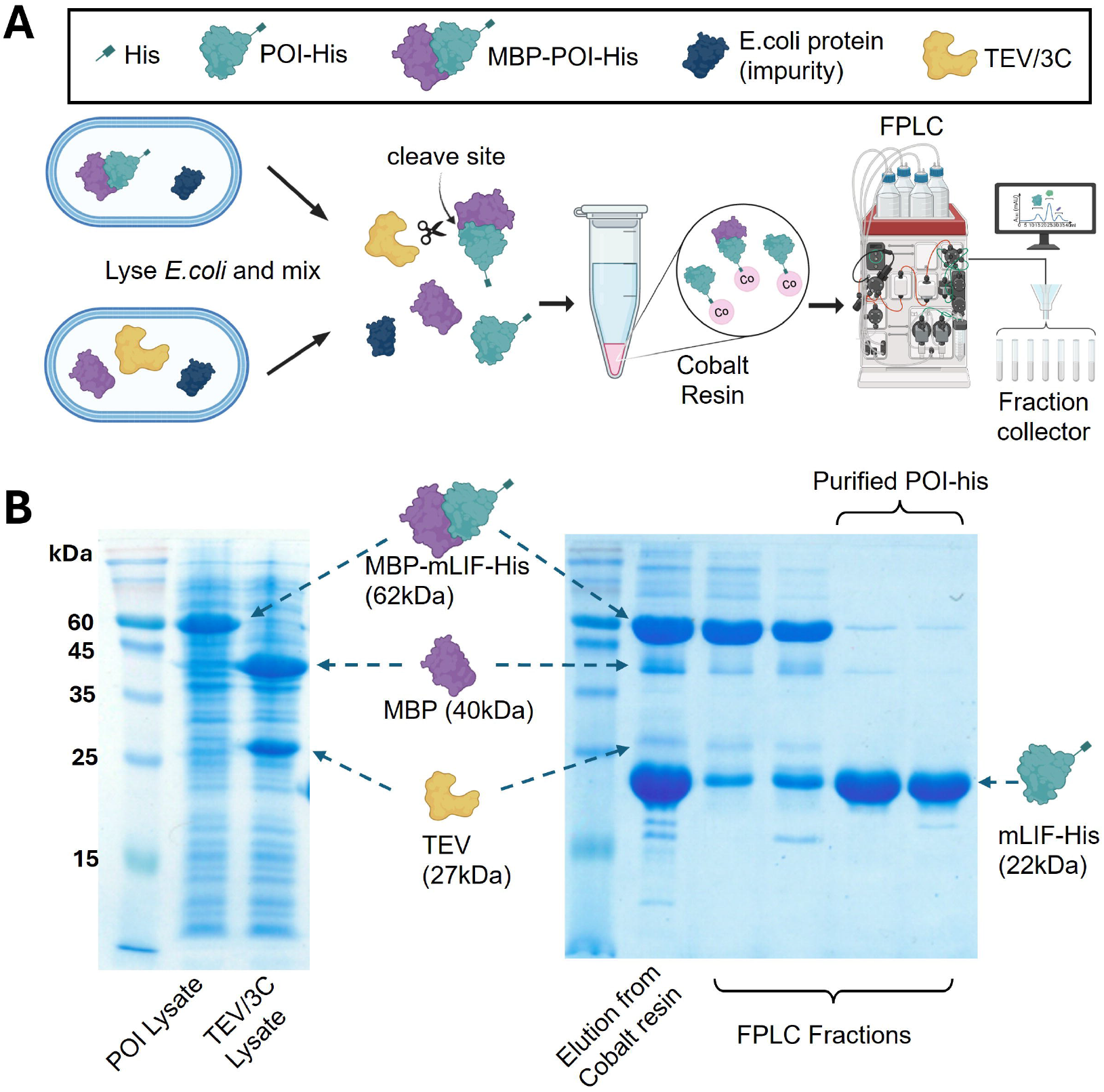
Overview of the protein purification workflow. (**A**) The protein of interest (POI) is expressed as a fusion construct consisting of an N-terminal maltose-binding protein (MBP) tag and a C-terminal histidine (His) tag, with a Tobacco etch virus (TEV) or human rhinovirus 3C protease cleavage site inserted between MBP and the POI. In parallel, a separate E. coli culture is used to express the corresponding solubility-tagged protease (MBP-TEV or MBP-3C). The two cultures are grown and induced independently, then lysed and combined at defined volume ratios (protease lysate:POI lysate = 1:7 for TEV and 1:15 for 3C). Upon mixing, the protease cleaves the MBP tag from the POI. Because the proteases have molecular weights close to the POI, excess fusion substrate is added to minimize protease carryover during downstream separation, as the uncleaved MBP-POI can be more efficiently separated from the cleaved POI by size-exclusion chromatography. Following immobilized metal ion affinity chromatography (IMAC), the elution is injected into fast protein liquid chromatography (FPLC) for final separation of the POI. (**B**) SDS-PAGE of samples collected at each step of the workflow, illustrated using mLIF purification as a representative example.

**Table 2.** Probe sonication parameters.

| Parameter | Value |
| --- | --- |
| Sonication ON | 2 s |
| Sonication OFF | 5 s |
| Power | 112.5 W |
| Time per cycle | 8 min |
| Total cycles (per 1 L culture) | 5 |

**Table 3.** SEC parameters for cytokine purification.

| Cytokines | Collection Window<br>1 column volume (CV) = 24 mL | Fraction Size | Elution Buffer | Fractions Collected |
| --- | --- | --- | --- | --- |
| mLIF | 0.32 CV - 0.72 CV | 0.50 mL | PBS | 8 - 9 |
| hIL-2 | 0.42 CV – 0.54 CV | 0.15 mL | PBS | 8 - 12 |
| mIL-33 | 0.36 CV - 0.48 CV | 0.15 mL | PBS | 8 - 11 |
| hIL-33 | 0.36 CV - 0.48 CV | 0.15 mL | PBS | 8 - 11 |

### Endotoxicity and Bioactivity

#### Determine the Endotoxicity Levels of Cytokines

**Timing: 3 days**

This step evaluates whether pRARE-containing ClearColi provides an endotoxin-free platform for recombinant protein production as demonstrated in **Figure S2**. Reagents used for this step are listed in Table S2.

1. Seed HEK-Blue™ hTLR4 cells into a 96-well plate at a density of 3 × 10^5^ cells/mL in 100 μL per well of Dulbecco’s Modified Eagle’s Medium (DMEM) supplemented with 10% fetal bovine serum (FBS) and 1% penicillin/streptomycin.
  a. Incubate cells overnight at 37°C with 5% CO_2_.
2. Aspirate the medium and replace with 100 μL per well of fresh DMEM supplemented with 10% heat inactivated FBS (56°C for 30 min) and 1% penicillin/streptomycin.
3. Treat each well with 10 μL of cytokine produced from pRARE-containing ClearColi™ or BL21 in a dilution series prepared in PBS.
  a. Begin with 1 μg/mL cytokine per well and perform 10-fold serial dilutions.
  b. Incubate cells overnight at 37°C with 5% CO_2_.
4. Transfer 20 μL of medium from each well into a new 96-well plate.
5. Add 180 μL of QUANTI-Blue™ assay buffer to each well.
  a. Incubate at 37°C and monitor every 20 min until a visible color difference is observed.
  b. Measure absorbance at 635 nm using a plate reader. **Note:** When the supernatant of HEK-Blue™ hTLR4 reporter cells is mixed with QuantiBlue substrate, the resulting blue color indicates the SEAP secretion from cells due to TLR4 recognizing endotoxin.

#### Determine the bioactivity of recombinant mLIF

**Timing: 2 weeks**

This step evaluates whether purified recombinant mLIF (**Figure 2A**) retains biological activity comparable to commercial standards. Reagents used for this step are listed in **Table S3**.

**Figure 2.**
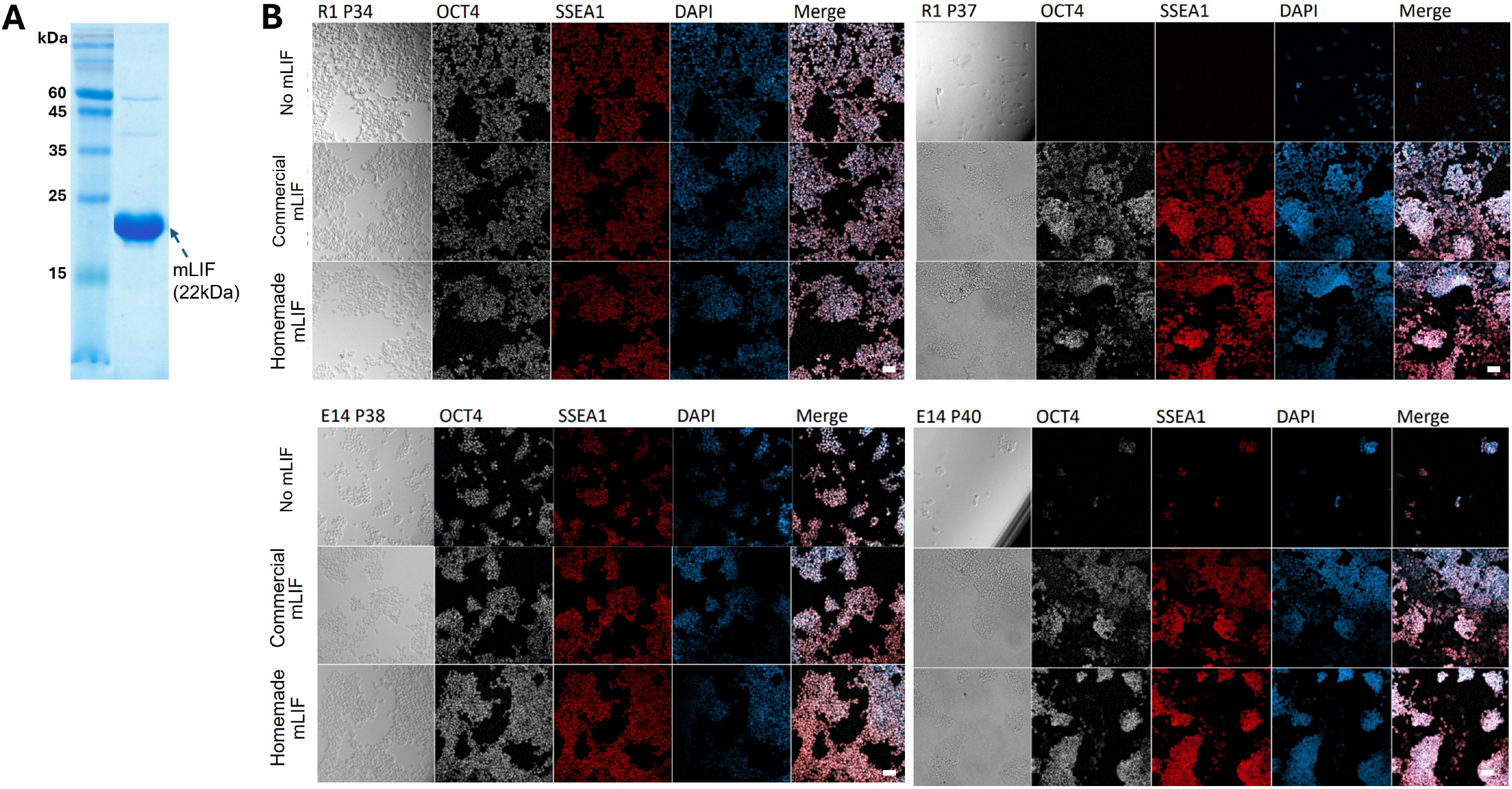
Purity and bioactivity validation of mLIF. (**A**) The purified homemade mLIF cytokine on a 15% SDS-PAGE gel. (**B**) Surface marker staining of SSEA1 and intracellular staining of OCT4 transcription factor show that both commercial and purified homemade mLIF have comparable bioactivity in maintaining mouse ESCs (lines R1 and E14) in its stem cell and undifferentiated state. Untreated mouse ESCs differentiate and decrease significantly in cell numbers with increasing passage counts. After each passage, cells were counted and seeded at the same density of 5 × 10 cells/cm^2^. DAPI was used for nuclear staining. Scale bar = 50 μm.

1. Coat 12-well tissue culture plates with 0.2% gelatin for 20 minutes.
  a. Air-dry the plates overnight.
2. Seed mouse naïve embryonic stem cells (ESCs) R1 cell line at a density of 5 × 10^4^ cells/cm^2^.
  a. Culture cells in either DMEM high-glucose medium supplemented with 15% ESC-qualified FBS, 1× GlutaMAX™, 1 mM sodium pyruvate, 1× MEM non-essential amino acids, and 1× 2-mercaptoethanol. **Note:** Alternatively, mouse ESCs E14 (ES-E14TG2A) cell line can be cultured in the same density in serum-free 2i medium consisting of a 1:1 ratio of DMEM/F12 to Neurobasal medium supplemented with 1× B-27 supplement, 0.5× N-2 supplement, 1× 2-mercaptoethanol, MEK inhibitor, and GSK3 inhibitor. Both R1 and E14 cell lines were used to evaluate mLIF bioactivity and confirm functionality across multiple mouse ESC lines.
3. Treat wells with either commercial mLIF or purified recombinant mLIF at a final concentration of 10 μg/mL. *Note:* Include untreated wells as a control group.
4. Incubate mouse ESCs at 37°C with 5% CO2.
5. Passage cells at approximately 80% confluency using TrypLE™ Express Enzyme.
  a. Gently rinse cells with prewarmed PBS and aspirate all remaining liquid.
  b. Add 300 μL TrypLE™ Express Enzyme and incubate at 37°C for 3 min.
  c. Quench the reaction with DMEM/F12 medium supplemented with 0.1% BSA.
  d. Centrifuge at 300 × g for 5 min.
  e. Discard the supernatant.
  f. Repeat steps c–e once.
  g. Resuspend the cells in cell culture media and seed the cells at a density of 5 × 10^4^ cells/cm^2^ in a new 12-well tissue culture plate.
6. Collect cells for immunostaining.
  a. Wash ESCs with PBS twice.
  b. Stain ESCs with 100 μL Stage-Specific Embryonic Antigen-1 (SSEA1) antibody at a 1:200 dilution in blocking buffer (PBS containing 0.1% Triton X-100 and 10% FBS) overnight at 4°C in the dark. i Wash ESCs twice with PBS supplemented with 0.05% Tween20.
  c. Fix cells with 100 μL fixation buffer for 10 minutes at room temperature.
  d. Permeabilize and block cells overnight at 4°C in the dark using blocking buffer.
  e. Stain ESCs with 100 μL rabbit anti-OCT4 antibody at a 1:200 dilution in blocking buffer overnight at 4°C in the dark. i Wash ESCs twice with PBS supplemented with 0.05% Tween20.
  f. Stain ESCs with 100 μL goat anti-rabbit IgG secondary antibody at a 1:1000 dilution overnight at 4°C in the dark. i Wash ESCs twice with PBS supplemented with 0.05% Tween20.
  g. Stain ESCs with DAPI at 0.1 μg/mL for 20 minutes at room temperature in the dark. i Wash ESCs twice with PBS supplemented with 0.05% Tween20.
  h. Observe fluorescence and cell morphology using a fluorescence microscope.
7. Repeat steps 1-6 at each passage until differences between mLIF-treated and untreated groups are observed.

**Note:** mLIF retains the undifferentiated state of mouse ESCs as demonstrated by OCT4 and SSEA1 staining (**Figure 2B**)^10^. In addition, homemade mLIF also maintained a similar cell morphology as compared to commercial mLIF for both E14 and R1 cell lines over the course of 5 days (**Figure S3**).

#### Determine the bioactivity of recombinant hIL-2

**Timing: 3 days**

This step evaluates whether purified recombinant hIL-2 (**Figure 3A**) retains biological activity comparable to commercial standards as illustrated in **Figure 3B**. Reagents used for this step are listed in **Table S4**.

**Figure 3.**
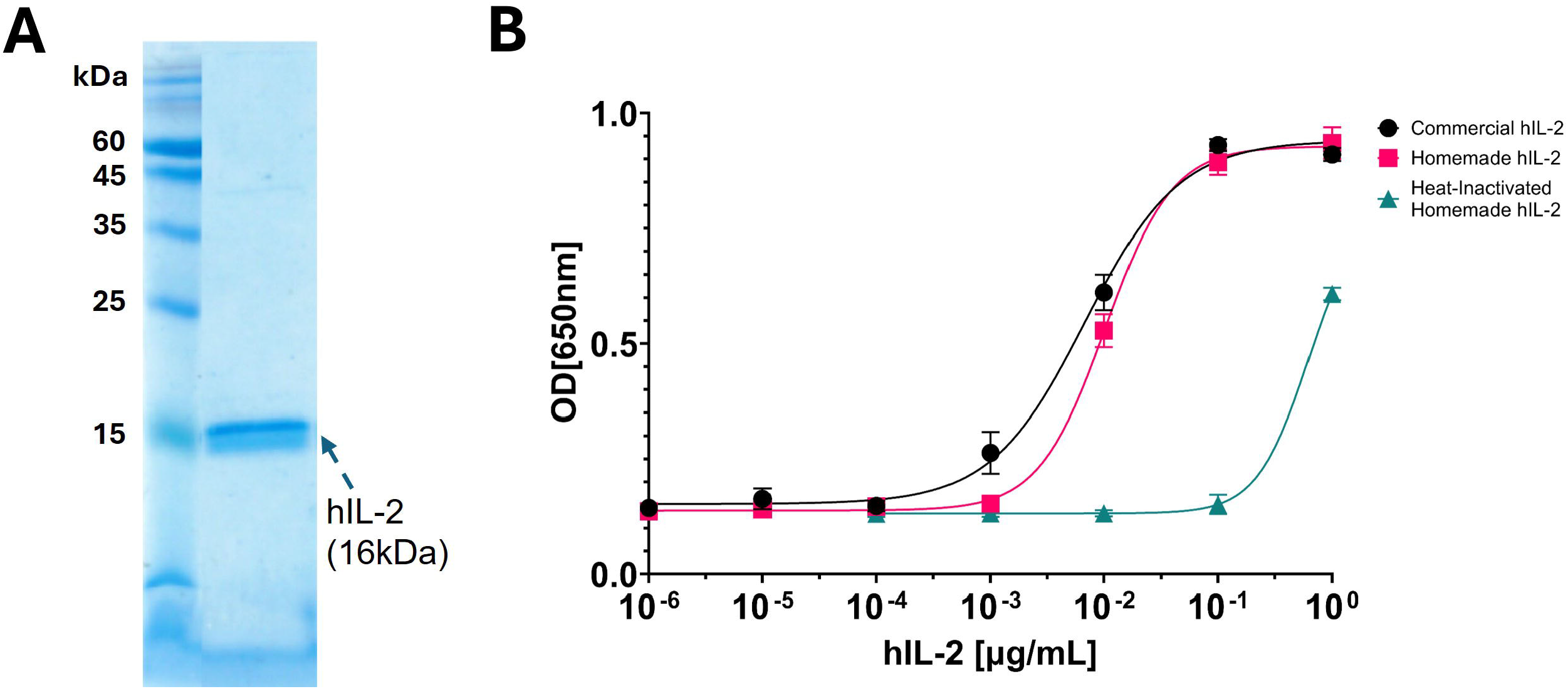
Purity and bioactivity validation of hIL-2. (**A**) The purified homemade hIL-2 cytokine on a 15% SDS-PAGE gel. **(B)** HEK-Blue^™^ hIL-2 reporter cells treated with commercial or purified homemade hIL-2 shows comparable bioactivity. Following stimulation, supernatants from reporter cells were collected and incubated with Quanti-Blue substrate, and the resulting colorimetric signal was quantified at 650 nm. Error bars represent standard deviations from 3 technical replicates (n = 3).

1. Seed HEK-Blue™ IL-2/IL-15 reporter cells into a 96-well plate at a density of 3 × 10 cells/mL in 100 μL per well of DMEM supplemented with 10% FBS and 1% penicillin/streptomycin.
  a. Incubate cells overnight at 37°C with 5% CO_2_.
2. Aspirate the medium and replace with 100 μL per well of fresh DMEM supplemented with 10% heat-inactivated FBS (56°C for 30 min) and 1% penicillin/streptomycin.
3. Treat each well with 10 μL of commercial hIL-2 or hIL-2 produced from pRARE-containing ClearColi in a dilution series prepared in PBS.
  a. Begin with 1 μg/mL hIL-2 per well and perform 10-fold serial dilutions.
  b. Incubate cells overnight at 37°C with 5% CO_2_. **Note:** Include heat-inactivated hIL-2 (95°C for 5 min) as a control group.
4. Transfer 20 μL of medium from each well into a new 96-well plate.
5. Add 180 μL of QUANTI-Blue™ assay buffer to each well.
  a. Incubate at 37°C and monitor every 20 min until a visible color difference is observed.
  b. Measure absorbance at 650 nm using a plate reader.

**Note:** When the supernatant of HEK-Blue™ IL-2 reporter cells is mixed with QuantiBlue substrate, the resulting blue color indicates the SEAP secretion from cells due to hIL-2 recognition.

Determine the bioactivity of recombinant mIL-33

**Timing: 4 days**

This step evaluates whether purified recombinant mIL-33 (**Figure 4A**) retains biological activity comparable to commercial standards. Reagents used for this step are listed in **Table S5**.

**Figure 4.**
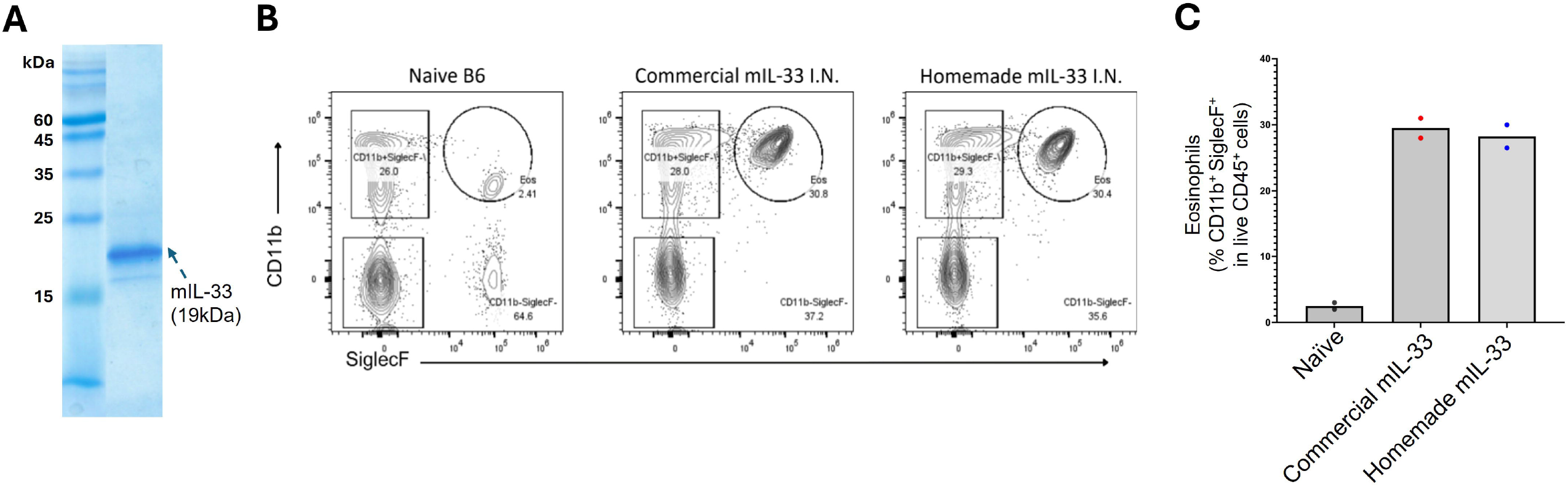
Purity and bioactivity validation of mIL-33. (**A**) The purified homemade mIL-33 cytokine on a 15% SDS-PAGE gel. (**B**) Gating strategy for eosinophil (FSC/SSC, FSC-A/FSC-H (Singlets), Viability (LD), CD45^+^, CD11b^+^ SiglecF^+^) in lung tissue of naïve B6 mice and B6 mice treated with commercial or purified homemade mIL-33. (**C**) Eosinophil percentage of live immune cells in B6 mouse lung tissue show comparable frequency between commercial and purified homemade mIL-33 treatment.

**Note:** This protocol is adapted from Kabil et al. for IL-33-induced allergic lung inflammation^11^.

1. Administer 0.5 μg of commercial mIL-33 or purified recombinant mIL-33 intranasally to adult 10-week-old C57BL/6J mice in 40 μL sterile PBS daily for 3 consecutive days. Administer PBS alone as a control.
  a. Euthanize mice and harvest lung tissues on day 4, 24 hr after the final challenge.
2. Mince lungs into small pieces and digest in RPMI 1640 supplemented with 10% FBS, 42.4 μg/mL Liberase, and 0.05 mg/mL DNase I for 40 min at 37°C in a rotating incubator.
3. Pass the digested tissue through a 40 μm cell strainer and wash with Fluorescence-Activated Cell Sorting (FACS) buffer (PBS containing 2% FBS, 2 mM EDTA, and 0.05% sodium azide)
  a. Centrifuge at 500 × g for 5 min.
  b. Discard supernatant.
4. Lyse red blood cells using 1 mL of ACK lysis buffer for 4 min at room temperature.
5. Quench samples with 20 mL FACS buffer.
  a. Centrifuge at 500 × g for 5 min.
  b. Discard supernatant.
  c. Resuspend pellet in 1 mL FACS buffer to obtain a single-cell suspension.
6. Incubate cells with LIVE/DEAD fixable dead cell stain using the LIVE/DEAD Fixable Near-IR Viability Kit according to the <u>manufacturer’s instructions</u>, together with TruStain FcX (anti-mouse CD16/32) antibody diluted 1:200 in PBS for 30 min at 4°C.
7. Incubate cells with fluorochrome-conjugated CD45, CD4, CD11b, CD11c, SiglecF antibodies for 30 min at 4°C.
8. Wash the cells with FACS buffer
  a. Centrifuge at 500 × g for 5 min.
  b. Discard supernatant.
  c. Resuspend pellet in 1 mL FACS buffer.
  d. Repeat step 8.
9. Analyze samples on a flow cytometer.

**Note:** Functional mIL-33 should induce eosinophilic inflammation in mouse lungs^11^ compared to naïve, PBS-treated mouse lungs as shown in **Figures 4B and 4C**.

#### Determine the bioactivity of recombinant hIL-33

**Timing: 3 days**

This step evaluates whether recombinant hIL-33 (**Figure 5A**) retains biological activity comparable to commercial standards. Reagents used for this step are listed in *Table S6*.

**Figure 5.**
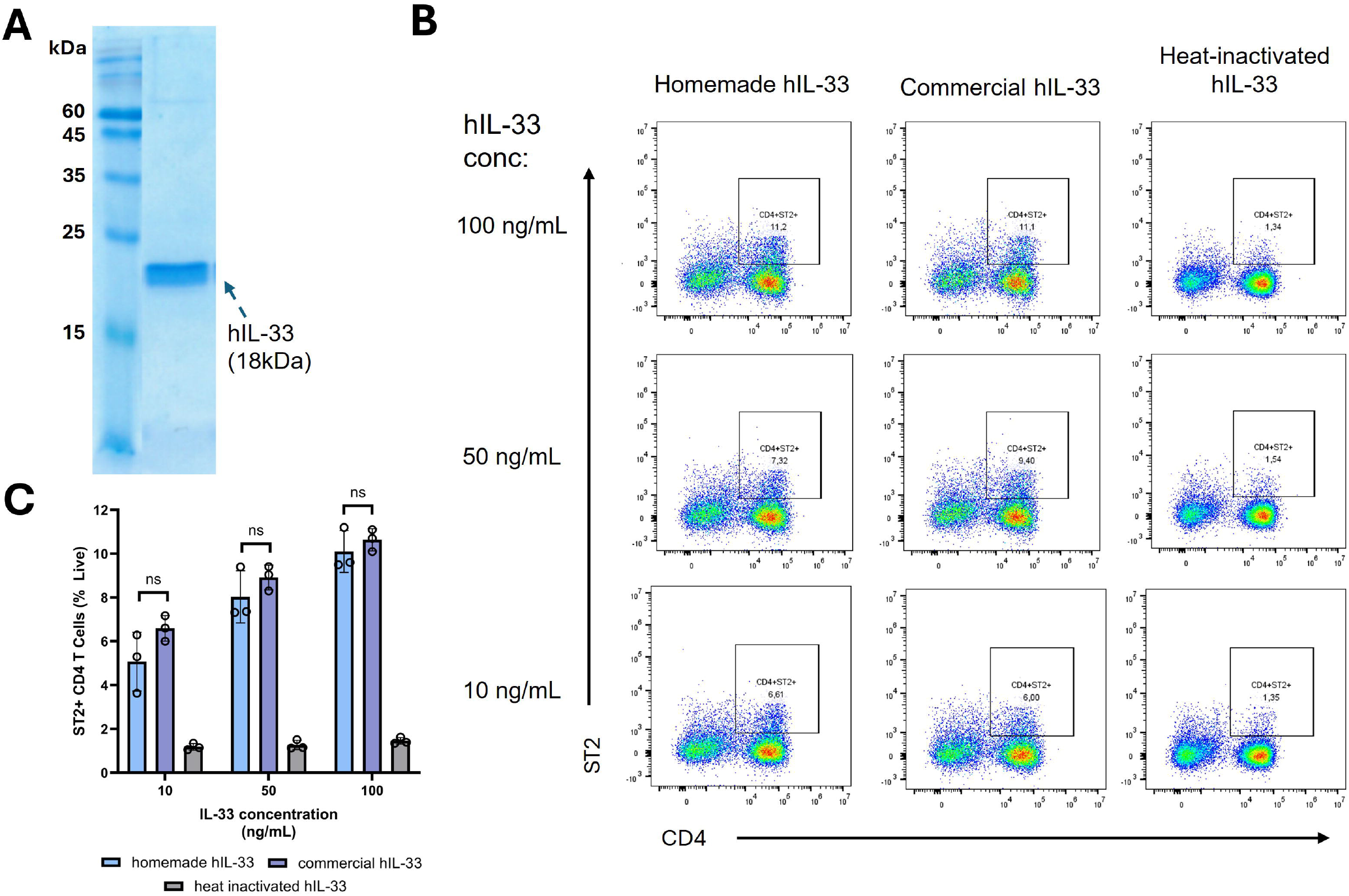
Purity and bioactivity validation of hIL-33. (**A**) The purified homemade hIL-33 cytokine on a 15% SDS-PAGE gel. (**B**) Gating strategy for T cells (FSC/SSC, FSC-A/FSC-H (Singlets), Viability (LD), CD45^+^, CD4^+^ ST2^+^) in peripheral blood mononuclear cells (PBMCs) treated with commercial or purified homemade hIL-33. (**C**) T cell (CD4^+^ ST2^+^) percentage of live PBMCs show comparable frequency between commercial and purified homemade hIL-33 treatment based on 2-way Anova multiple comparisons.

1. Prepare 20 mL of thawing media consisting of ImmunoCult-XF expansion medium supplemented with 20 ng/mL IL-2.
2. Thaw cryopreserved primary human peripheral blood mononuclear cells (PBMCs) in a 37°C water bath into prepared thawing media.
  a. Centrifuge the cells at 500 x g for 5 minutes.
  **Note:** Cryopreserved PBMCs were used for bioactivity validation as they were readily available. Alternatively, freshly prepared PBMCs can be used to evaluate hIL-33 bioactivity.
3. Negatively isolate T cells using EasySep^™^ Human T Cell Isolation Kit as per <u>manufacturer’s instructions</u>.
4. Resuspend the final T cell pellet in ImmunoCult medium supplemented with 20 ng/mL IL-2 and 25 μL/mL CD3/CD28 activator.
5. Seed 1 × 105 cells per well in a 96-well plate.
6. Add commercial hIL-33 or purified recombinant hIL-33 to designated wells at final concentrations of 10, 50, or 100 ng/mL.
  a. Incubate cells for 3 days at 37°C with 5% CO_2_. **Note:** Include heat-inactivated hIL-33 (98°C for 15 min) as a control group.
7. Monitor T cell activation by assessing cell clustering under a microscope.
8. Analyze cellular phenotype for ST2-expressing CD4 T cells using flow cytometry on day 3 of activation.

**Note:** Functional hIL-33 should bind to the ST2 receptor and activate ST2-expressing CD4^+^ T cells isolated from human PBMCs^12^ as shown in **Figures 5B and 5C**.

## Expected outcomes

Successful protein purification from a 4 L bacterial culture is expected to yield 0.8–3.6 mg of high-purity, functionally active protein, depending on the specific cytokine as shown in Table 4.Recombinant mLIF, mIL-33, and hIL-33 typically yield approximately 0.8 mg of protein and can be detected as bands at approximately 20 kDa (**Figure 2A**), 17.5 kDa (**Figure 4A**), and 17.5 kDa (**Figure 5A**), respectively, on SDS-PAGE gels. Recombinant hIL-2 yields approximately 3.6 mg of protein and is detected as an approximately 16.7 kDa band (**Figure 3A**). Uncropped SDS-PAGE gels for these figures are presented in **Figure S4**.

**Table 4.**
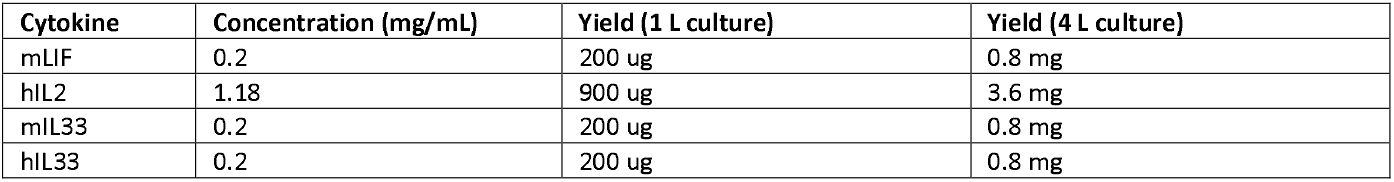
Cytokine yield and concentration following purification.

## Limitations

This protocol provides an accessible and user-friendly approach for cytokine production, enabling research labs and core facilities to scale protein expression and purification. However, one limitation should be considered. The protocol requires access to fast protein liquid chromatography (FPLC) to achieve high-purity cytokine isolation. With only IMAC, non-specific binding of E. coli impurity proteins, MBP, and proteases lacking His tags can occur, leading to their presence in the elution. Use of alternative FPLC systems or columns may require optimization of elution parameters, resulting in variability between setups.

## Troubleshooting

### Problem 1

No bacterial growth.

#### Potential solution

Ensure that the correct antibiotics and working concentrations are used for the bacterial strain and plasmid selection. Incorrect antibiotic selection or concentration can prevent growth of transformed cells. Verify plasmid resistance markers and prepare fresh antibiotic stocks if necessary.

### Problem 2

High levels of impurities observed on SDS-PAGE gels.

#### Potential solution

Non-specific binding of E. coli proteins to the HisPur− cobalt resin may result in co-elution with the target cytokine. To reduce nonspecific binding, increase the imidazole concentration in the protein binding buffer to 10-30 mM.

#### CRITICAL

Excessively high imidazole concentrations may prematurely elute the target protein, leading to reduced cytokine yield. Optimization within this range is recommended.

### Problem 3

Cytokines are not detected on SDS-PAGE gels.

#### Potential solution

Protein loss may occur at multiple stages of the purification workflow. To identify the source of loss, analyze samples from each step of the protocol by SDS-PAGE. First, if the protein is not detected in the total bacterial lysate, protein expression may not have been successfully induced. Confirm that the cytokine plasmid contains an appropriate inducible promoter (e.g., lac operon) and that IPTG induction conditions are correct. On the other hand, if the protein is detected in the pellet after sonication but not in the soluble lysate, the cytokine is likely insoluble. Confirm that the construct includes a solubility-enhancing tag (e.g., MBP). Finally, if the protein is present in the lysate but does not bind to the HisPur ^−^cobalt resin, the 6×His tag may be inaccessible. Consider introducing flexible linkers between the cytokine and the His tag to improve binding efficiency.

### Problem 4

Cytokines have suboptimal bioactivity compared to commercial standards.

#### Potential solution

Protein stability may decrease following repeated freeze-thaw cycles, leading to reduced bioactivity. To maintain optimal bioactivity, store proteins in aliquots and thaw only the amount required immediately before use.

## Resource availability

### Lead contact

For further information or requests for resources and reagents, please contact the lead author, Yanpu He.

### Materials availability

All plasmids generated for protein purification in this study are available from the lead contact.

## Acknowledgments

This work was supported by startup funds from the School of Biomedical Engineering at the University of British Columbia to YH. YH is a Canada Research Chair (Tier 2)

KMM is supported by a Canadian Stem Cell Network Impact Award (IMP-C5R1-5) and a CIHR Project Grant (PJT186035). ICK is a trainee in the KMM laboratory and is supported by the Killam Foundation Doctoral Award and the NSERC CREATE ImmunoEngineering Doctoral Scholarship.

NS is supported by the Natural Sciences and Engineering Research Council of Canada (NSERC, award no. RGPIN-2020-04198), a Canadian Institutes of Health Research Project Grant, and startup funds from the School of Biomedical Engineering at the University of British Columbia. NS is the recipient of a Michael Smith Health Research BC Scholar Award and a Canada Research Chair (Tier 2).

## Author contributions

JYK and YH designed the purification approach, cloned the expression plasmids, and optimized the purification procedures, with JX helping with protein purification. JYK validated the activity of hIL-2. ICK and SXC validated the activity of hIL-33. AK validated the activity of mIL-33. KWC validated the activity of mLIF. YH, KMM, and NS supervised the experiments. JYK wrote the paper with revisions from all authors.

## Declaration of interests

The authors have no conflicts of interest to declare.

---

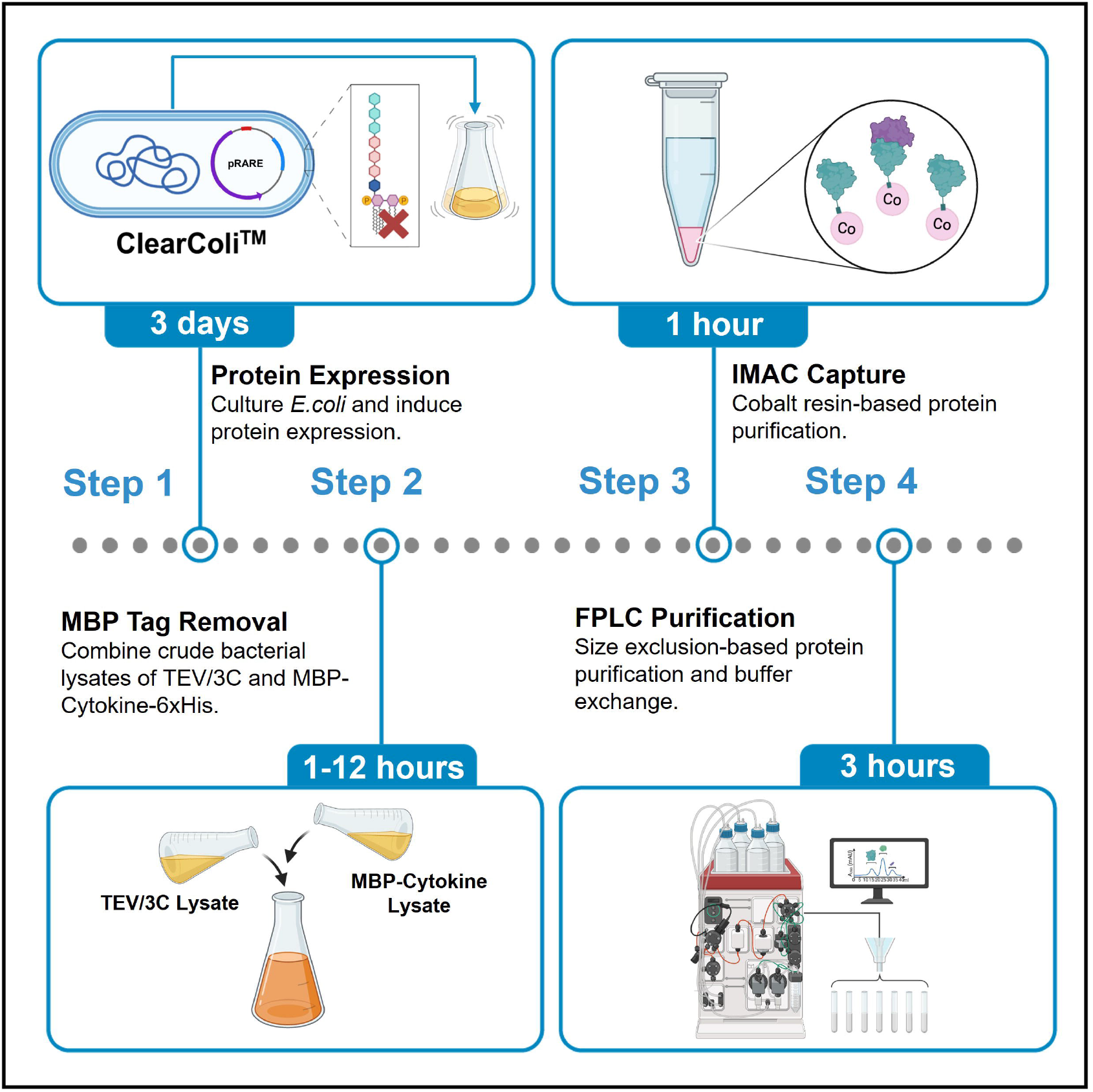

## Notes

### Competing Interest Statement

The authors have declared no competing interest.

## References

1. Kungwankiattichai, S., and Maziarz, R.T. (2025). The history of cytokines and growth factors development. Best Practice & Research Clinical Haematology 38, 101612. 10.1016/j.beha.2025.101612.

2. Mamat, U., Wilke, K., Bramhill, D., Schromm, A.B., Lindner, B., Kohl, T.A., Corchero, J.L., Villaverde, A., Schaffer, L., Head, S.R., et al. (2015). Detoxifying Escherichia coli for endotoxin-free production of recombinant proteins. Microb Cell Fact 14, 57. 10.1186/s12934-015-0241-5.

3. Mamat, U., Woodard, R.W., Wilke, K., Souvignier, C., Mead, D., Steinmetz, E., Terry, K., Kovacich, C., Zegers, A., and Knox, C. (2013). Endotoxin-free protein production—ClearColi technology. Nat Methods 10, 916–916. 10.1038/nmeth.f.367.

4. Wilding, K.M., Hunt, J.P., Wilkerson, J.W., Funk, P.J., Swensen, R.L., Carver, W.C., Christian, M.L., and Bundy, B.C. (2019). Endotoxin-Free E. coli-Based Cell-Free Protein Synthesis: Pre-Expression Endotoxin Removal Approaches for on-Demand Cancer Therapeutic Production. Biotechnol. J. 14, 1800271. 10.1002/biot.201800271.

5. Tegel, H., Tourle, S., Ottosson, J., and Persson, A. (2010). Increased levels of recombinant human proteins with the Escherichia coli strain Rosetta(DE3). Protein Expression and Purification 69, 159–167. 10.1016/j.pep.2009.08.017.

6. Guo, Y., Yu, M., Jing, N., and Zhang, S. (2018). Production of soluble bioactive mouse leukemia inhibitory factor from Escherichia coli using MBP tag. Protein Expression and Purification 150, 86–91. 10.1016/j.pep.2018.05.006.

7. LGC 60810, ClearColi BL21(DE3) Electrocompetent Cells, 12 Reactions (DUOs) https://www.sigmaaldrich.com/CA/en/product/sigma/lgc608101?srsltid=AfmBOop6qh3zCv0j2iBcIiH4Lf2vUWvmsBJTJOiaLF1bGWvcNLlxuBeG.

8. Green, M.R., and Sambrook, J. (2020). The Inoue Method for Preparation and Transformation of Competent Escherichia coli: “Ultracompetent” Cells. Cold Spring Harb Protoc 2020, 101196. 10.1101/pdb.prot101196.

9. Raran-Kurussi, S., and Waugh, D.S. (2016). A dual protease approach for expression and affinity purification of recombinant proteins. Analytical Biochemistry 504, 30–37. 10.1016/j.ab.2016.04.006.

10. Hirai, H., Firpo, M., and Kikyo, N. (2015). Establishment of leukemia inhibitory factor (LIF)-independent iPS cells with potentiated Oct4. Stem Cell Research 15, 469–480. 10.1016/j.scr.2015.09.002.

11. Kabil, A., Nayyar, N., Brassard, J., Li, Y., Chopra, S., Hughes, M.R., and McNagny, K.M. (2024). Microbial intestinal dysbiosis drives long-term allergic susceptibility by sculpting an ILC2–B1 cell–innate IgE axis. Journal of Allergy and Clinical Immunology 154, 1260-1276.e9. 10.1016/j.jaci.2024.07.023.

12. Endo, Y., Hirahara, K., Iinuma, T., Shinoda, K., Tumes, D.J., Asou, H.K., Matsugae, N., Obata-Ninomiya, K., Yamamoto, H., Motohashi, S., et al. (2015). The Interleukin-33-p38 Kinase Axis Confers Memory T Helper 2 Cell Pathogenicity in the Airway. Immunity 42, 294–308. 10.1016/j.immuni.2015.01.016.

