## Supplemental information for "Cost-Effective Purification of Endotoxin-Free LIF, IL-2, and IL-33"

Figures S1-S4, Tables S1-S6.

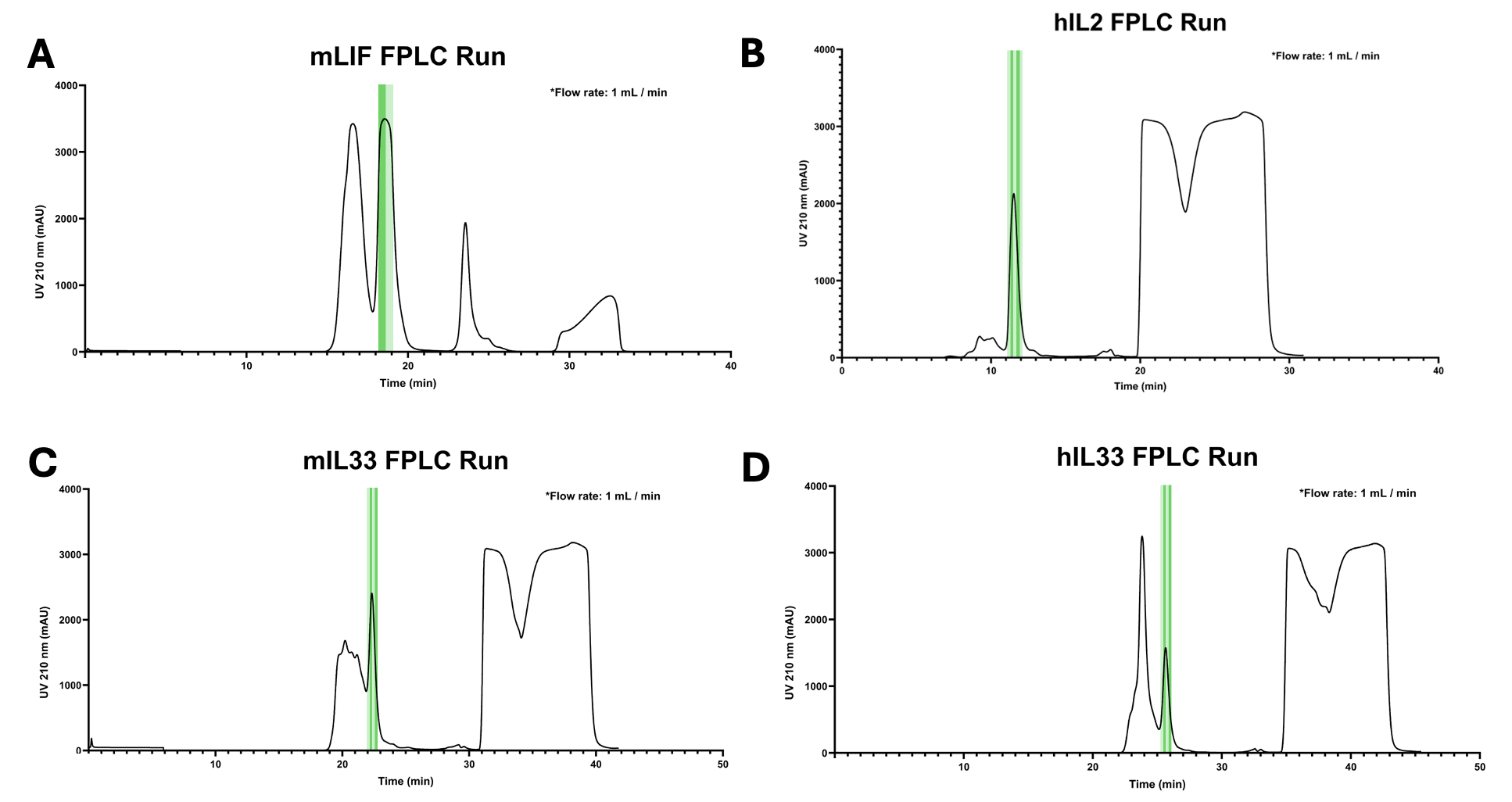

**Figure S1. Fast protein liquid chromatography plots of each cytokine**. Collected fractions are highlighted in green. Absorbance at 210 nm vs time plot of (A) mLIF, (B) hIL-2, (C) mIL-33, and (D) hIL-33 during FPLC runs.

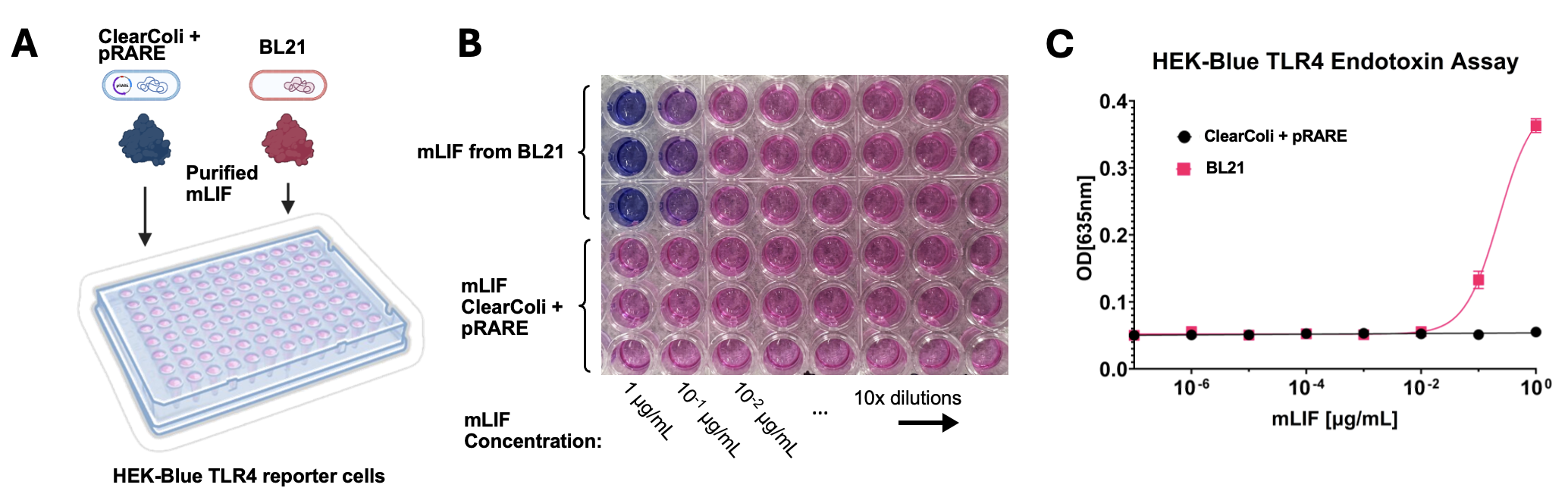

**Figure S2. Endotoxin assay shows that the protein production platform is endotoxin free.** (**A**) Schematic illustration of HEK-Blue™ TLR4 reporter cells treated with protein expressed from ClearColi with pRARE or BL21. (**B**) 12 hours post treatment, supernatant of reporter cells were added to a new plate and mixed with QuantiBlue substrate. Blue color indicate the SEAP secretion from cells due to TLR4 recognizing endotoxin. **(C**) Absorbance at 635 nm of QuantiBlue assay shows ClearColi-derived mLIF contains undetectable endotoxin up to 1 μg/mL.

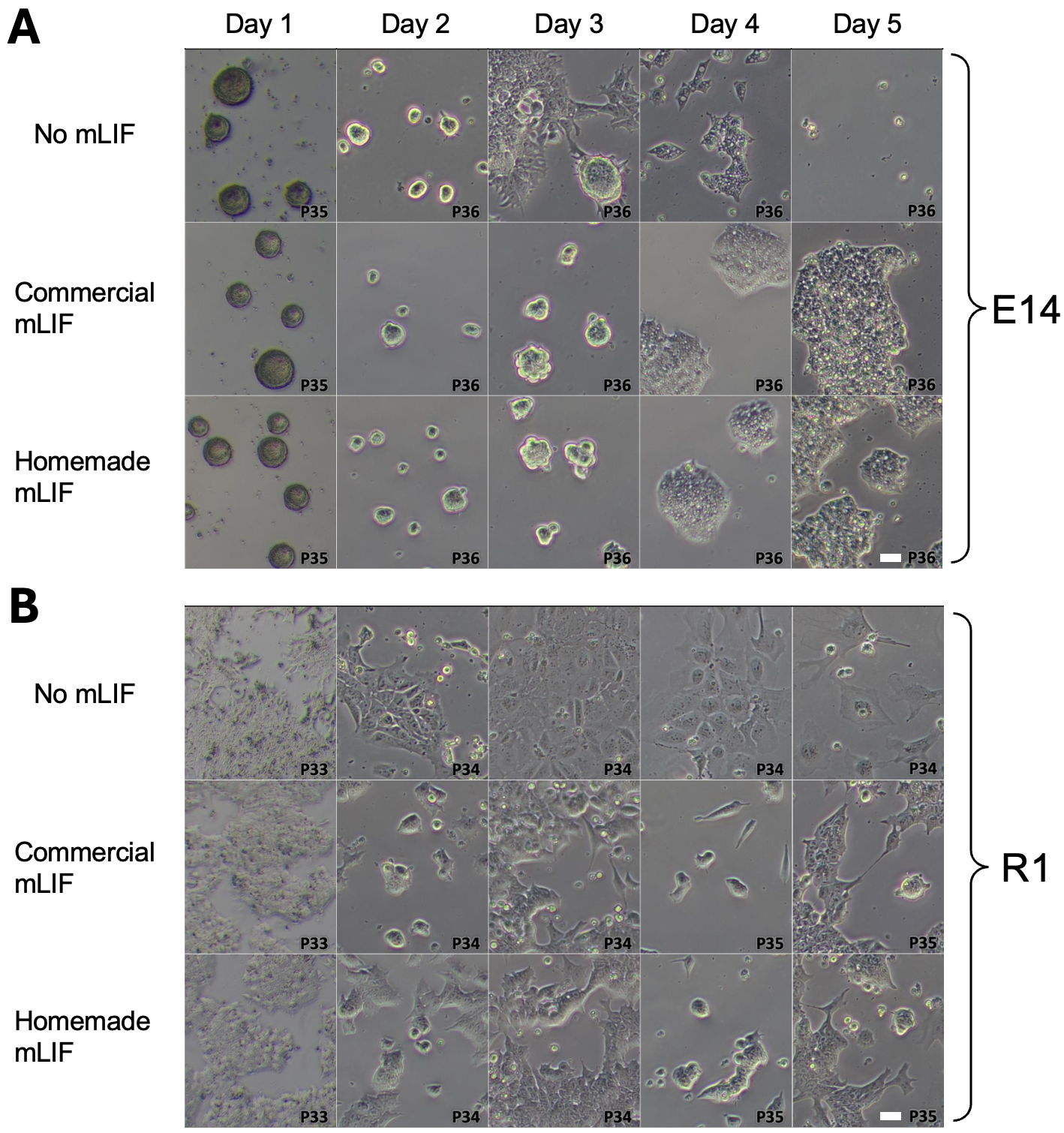

**Figure S3: Brightfield microscopy images showing the morphology** of (A) E14 and (B) R1 mouse embryonic stem cell lines treated with 10 μg/mL of commercial versus homemade mLIF over the course of 5 days, P indicates the passage number, scale bar = 20 μm.

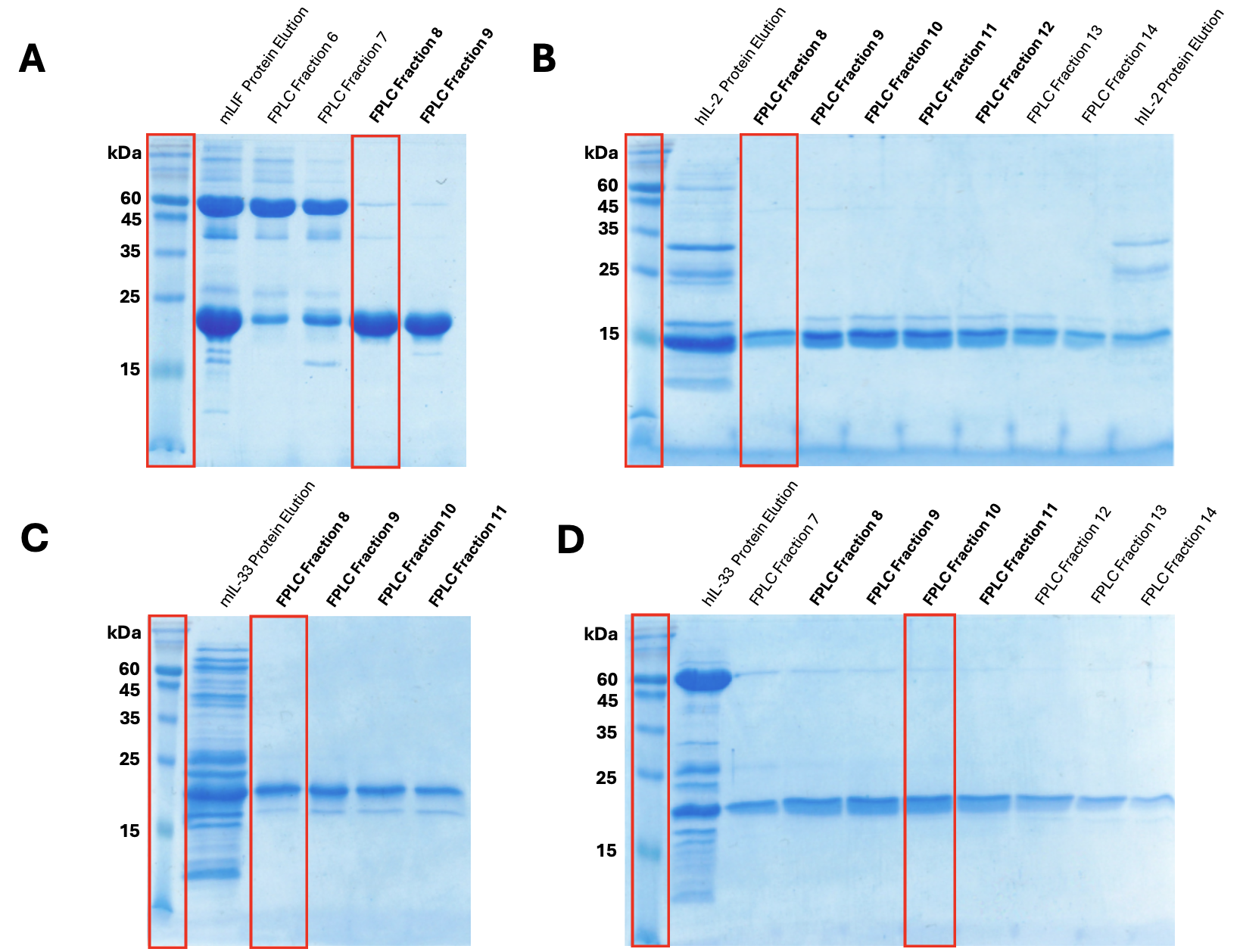

**Figure S4. Uncropped gel images of cytokine elution and the collected fractions from FPLC size-exclusion chromatography.** Outlined in red are gel sections showed in Figure 2, 3, 4 and 5. (A) Full gel image for mLIF. (B) Full gel image for hIL-2. (C) Full gel image for mIL-33. (D) Full gel image for hIL-33.

Table S1: Sequences for cytokine plasmid design.

| **COMPONENT** | **SEQUENCE** |
| --- | --- |
| Tac promoter | ttgacaattaatcatcggctcgtataatg |
| Lac operator | ttgtgagcggataacaa |
| MBP-tag | atgaaaatcgaagaaggtaaactggtaatctggattaacggcgataaaggctataacggtctcgctgaagtcggtaagaaattcgagaaagataccggaattaaagtcaccgttgagcatccggataaactggaagagaaattcccacaggttgcggcaactggcgatggccctgacattatcttctgggcacacgaccgctttggtggctacgctcaatctggcctgttggctgaaatcaccccggacaaagcgttccaggacaagctgtatccgtttacctgggatgccgtacgttataacggcaaactgattgcttaccccatcgctgttgaagcgttatcgctgatttataacaaagatcttctgccgaacccgccaaaaacctgggaagagatcccggcgctggataaagaactgaaagcgaaaggtaagagcgcgctgatgttcaacctgcaagaaccgtacttcacctggccgctgattgctgctgacgggggttatgcgttcaagtatgaaaacggcaagtacgacattaaagacgtgggcgtggataacgctggcgcgaaagcgggtctgaccttcctggttgacctgattaaaaacaaacacatgaatgcagacaccgattactccatcgcagaagctgcctttaataaaggcgaaacagcgatgaccatcaacggcccgtgggcatggtccaacatcgacaccagcaaagtgaattatggtgtaacggtactgccgaccttcaagggtcaaccatccaaaccgttcgttggcgtgctgagcgcaggtattaacgccgccagtccgaacaaagagctggcaaaagagttcctcgaaaactatctgctgactgatgaaggtctggaagcggttaataaagacaaaccgctgggtgccgtagcgctgaagtcttacgaggaagagttggcgaaagatccacgtattgccgccaccatggaaaacgcccagaaaggtgaaatcatgccgaacatcccgcagatgtccgctttctggtatgccgtgcgtactgcggtgatcaacgccgccagcggtcgtcagactgtcgatgaagccctgaaagacgcgcagactggatcc |
| Protease cut site | TEV: gagaatctgtacttccagggt  HRV 3C: ctggaagttctgttccaggggccc |
| Cytokine gene fragments | mLIF: CCCCTGCCGATAACCCCTGTTAATGCAACATGCGCCATTCGTCATCCGTGTCACGGAAATTTGATGAATCAGATTAAAAATCAGCTGGCGCAACTTAACGGCTCTGCCAACGCTCTGTTCATCAGCTATTATACCGCCCAAGGCGAACCGTTTCCGAACAACGTTGAGAAACTGTGTGCGCCAAATATGACTGATTTTCCTTCATTCCATGGTAACGGGACAGAGAAGACTAAACTTGTAGAATTATACCGCATGGTCGCTTATCTCTCGGCGAGCCTGACGAATATTACGCGTGATCAGAAGGTGCTGAATCCCACCGCGGTGTCCCTGCAGGTCAAACTGAACGCAACCATCGATGTTATGCGCGGCCTCTTAAGCAACGTGCTGTGTCGGTTGTGCAACAAATACCGCGTCGGTCACGTGGACGTCCCACCGGTTCCGGACCATAGTGATAAAGAAGCATTTCAACGTAAAAAACTGGGTTGCCAGTTACTAGGCACGTACAAGCAGGTAATCAGTGTGGTGGTACAGGccttcGGTAGCTCT  hIL-2:  ATGGCCCCAACGAGCAGTAGCACCAAGAAGACCCAGCTCCAACTGGAGCATCTGCTATTGGATTTGCAGATGATTCTCAATGGTATCAACAATTATAAAAACCCTAAACTTACCCGCATGCTGACGTTCAAATTCTATATGCCGAAAAAAGCGACTGAATTAAAACACCTGCAGTGTCTTGAGGAAGAATTAAAGCCCTTGGAAGAAGTTCTGAATCTGGCGCAATCCAAAAACTTCCATCTGCGTCCGCGGGACCTGATCTCTAACATCAATGTCATTGTACTGGAGCTGAAAGGCAGTGAAACGACATTTATGTGCGAATACGCAGATGAGACTGCTACCATAGTGGAATTTCTGAACCGCTGGATTACATTTGCCCAGTCGATTATCTCAACCTTAACG  mIL-33 (cleaved):  AGCATTCAGGGCACGTCTCTCTTGACCCAGTCGCCGGCAAGCTTAAGTACATACAACGATCAGTCAGTTTCGTTCGTACTGGAAAACGGCTGTTACGTTATCAATGTGGACGACTCAGGTAAGGATCAAGAGCAGGATCAAGTGCTGTTACGTTATTACGAAAGCCCATGTCCTGCTAGCCAATCAGGTGATGGTGTGGATGGCAAAAAACTGATGGTTAATATGAGTCCGATTAAAGATACTGATATCTGGCTACACGCCAATGATAAGGATTATTCGGTAGAACTGCAGCGCGGCGACGTGTCCCCGCCCGAGCAGGCGTTTTTTGTCTTGCATAAAAAAAGCTCCGACTTCGTCTCCTTTGAATGCAAAAACCTGCCGGGGACCTATATTGGAGTCAAGGACAACCAGCTGGCGCTTGTGGAAGAGAAAGACGAATCTTGCAATAACATAATGTTCAAACTGAGTAAAATC  hIL-33 (cleaved):  agtatcacaggaatttcacctattacagagtatcttgcttctctaagcacatacaatgatcaatccattacttttgctttggaggatgaaagttatgagatatatgttgaagacttgaaaaaagatgaaaagaaagataaggtgttactgagttactatgagtctcaacacccctcaaatgaatcaggtgacggtgttgatggtaagatgttaatggtaaccctgagtcctacaaaagacttctggttgcatgccaacaacaaggaacactctgtggagctccataagtgtgaaaaaccactgccagaccaggccttctttgtccttcataatatgcactccaactgtgtttcatttgaatgcaagactgatcctggagtgtttataggtgtaaaggataatcatcttgctctgattaaagtagactcttctgagaatttgtgtactgaaaatatcttgtttaagctctctgaaact |
| 6x His tag | catcatcaccatcaccat |

Table S2: Key resources for determining endotoxin levels.

| **REAGENT or RESOURCE** | **SOURCE** | **CATALOG #** |
| --- | --- | --- |
| HEK-Blue^TM^ hTLR4 cell line | InvivoGen | hkb-htlr4 |
| QUANTI-Blue^TM^ Solution | InvivoGen | rep-qbs |

Table S3: Key resources for testing the bioactivity of mLIF.

| **REAGENT or RESOURCE** | **SOURCE** | **CATALOG #** |
| --- | --- | --- |
| Gelatin | Millipore Sigma | G1890 |
| Mouse embryonic stem cell line R1 | ATCC | SCRC-1011 |
| Mouse embryonic stem cell line ES-E14TG2a | ATCC | CRL-1821 |
| Dulbecco’s Modified Eagle’s Medium (DMEM) - high glucose | Thermo Fisher Scientific | 11965092 |
| Embryonic stem cell-quality fetal bovine serum | Wisent | 920-040 |
| GlutaMAX^TM^ | Thermo Fisher Scientific | 35050061 |
| Sodium pyruvate | Thermo Fisher Scientific | 11360070 |
| MEM Non-essential amino acids solution | Thermo Fisher Scientific | 11140050 |
| 2-mercaptoethanol | Thermo Fisher Scientific | 21985023 |
| F12 medium | Thermo Fisher Scientific | 11330057 |
| Neurobasal medium | Thermo Fisher Scientific | 21103049 |
| B27 supplement | Thermo Fisher Scientific | 17504044 |
| N2 supplement | Thermo Fisher Scientific | 17502048 |
| MEK inhibitor (PD0325901) | Tocris Bioscience | 4192/10 |
| GSK3 inhibitor (CHIR99021) | Millipore Sigma | SML1046 |
| Recombinant mouse LIF | R&D Systems | 8878-LF |
| TrypLE^TM^ Express Enzyme | Thermo Fisher Scientific | 12604021 |
| Alexa Fluor^©^ 647 Mouse Anti-SSEA-1 antibody | BD Bioscience | 562277 |
| Fixation buffer | BioLegend | 420801 |
| Triton^TM^ X-100 | Millipore Sigma | X100 |
| Rabbit Anti-OCT4 antibody | Abcam | ab137427 |
| Alexa Fluor^©^ 594 Goat Anti-Rabbit IgG secondary antibody | Thermo Fisher Scientific | A-11012 |
| DAPI | Thermo Fisher Scientific | D3571 |
| Dulbecco’s Phosphate Buffered Saline (calcium and magnesium free) | Thermo Fisher Scientific | 14190144 |

Table S4: Key resources for testing the bioactivity of hIL-2.

| **REAGENT or RESOURCE** | **SOURCE** | **CATALOG #** |
| --- | --- | --- |
| HEK-Blue^TM^ IL-2/IL-15 cell line | InvivoGen | hkb-il2-2 |
| QUANTI-Blue^TM^ Solution | InvivoGen | rep-qbs |
| Human Recombinant IL-2 (CHO-expressed) | STEMCELL Technologies | 78036 |

Table S5: Key resources for testing the bioactivity of mIL-33.

| **REAGENT or RESOURCE** | **SOURCE** | **CATALOG #** |
| --- | --- | --- |
| Mouse Recombinant IL-33 | BioLegend | 580506 |
| C57BL/6J mice | The Jackson Laboratory | #000664 |
| RPMI 1640 medium | Thermo Fisher Scientific | 11875093 |
| Liberase^TM^ Research Grade | Roche | 5401127001 |
| DNase I | Roche | 10104159001 |
| ACK lysis buffer | Thermo Fisher Scientific | A1094201 |
| LIVE/DEAD^TM^ Fixable Near-IR \| Dead Cell Stain Kit | Thermo Fisher Scientific | L10119 |
| TruStain FcX^TM^ (anti-mouse CD16/32) | BioLegend | 101320 |
| BD Pharmingen^TM^ PE Rat Anti-Mouse Siglec-F antibody | BD Biosciences | 552126 |
| CD4 monoclonal antibody (GK1.5) | Thermo Fisher Scientific | 14-0041-82 |
| Alexa Fluor^©^ 700 anti-mouse CD45 antibody | BioLegend | 103128 |
| PE-Fire 700 anti-mouse CD11b antibody | BioLegend | 101288 |
| BUV737 anti-mouse CD11c antibody | BioLegend | 569236 |

Table S6: Key resources for testing the bioactivity of hIL-33.

| **REAGENT or RESOURCE** | **SOURCE** | **CATALOG #** |
| --- | --- | --- |
| ImmunoCult-XF T cell expansion media | STEMCELL Technologies | 10981 |
| Human Recombinant IL-2 (CHO-expressed) | STEMCELL Technologies | 78036 |
| EasySep^TM^ Human T Cell Isolation Kit | STEMCELL Technologies | 100-0695 |
| ImmunoCult Activator CD3/CD28 | STEMCELL Technologies | 10971 |
| Recombinant Human IL-33 (carrier-free) | BioLegend | 581804 |
